# A Biologically Plausible Spiking Neural Network for Decoding Kinematics in the Hippocampus and Premotor Cortex

**DOI:** 10.1101/2022.11.09.515838

**Authors:** Elijah Taeckens, Ryan Dong, Sahil Shah

## Abstract

This work presents a spiking neural network for predicting kinematics from neural data towards accurate and energy-efficient brain machine interface. A brain machine interface is a technological system that interprets neural signals to allow motor impaired patients to control prosthetic devices. Spiking neural networks have the potential to improve brain machine interface technology due to their low power cost and close similarity to biological neural structures. The SNN in this study uses the leaky integrate-and-fire model to simulate the behavior of neurons, and learns using a local learning method that uses surrogate gradient to learn the parameters of the network. The network implements a novel continuous time output encoding scheme that allows for regression-based learning. The SNN is trained and tested offline on neural and kinematic data recorded from the premotor cortex of a primate and the hippocampus of a rat. The model is evaluated by finding the correlation between the predicted kinematic data and true kinematic data, and achieves peak Pearson Correlation Coefficients of 0.77 for the premotor cortex recordings and 0.80 for the hippocampus recordings. The accuracy of the model is benchmarked against a Kalman filter decoder and a LSTM network, as well as a spiking neural network trained with backpropagation to compare the effects of local learning.

## I. Introduction

Each year, there are about 18,000 new cases of spinal cord injuries in the United States [1]. These injuries frequently lead to severe degradation of motor function. Recent advancements have allowed the development of Brain Machine Interfaces (BMI), which can help restore patient mobility by decoding neural signals and using them to control prosthetic devices [2] [3]. This technology can greatly improve quality of life by improving patient mobility and freedom.

BMI are typically classified into invasive methods and non-invasive methods. While non-invasive methods, such as electroencephalography, have had some success with simple tasks, they lack the spatial and temporal resolution of intracortical recordings [4]. Classically, the most successful intracortical BMI have been based on applying Kalman filters or similar linear filters to decode neural signals [5]. More recent attempts have also proven that machine learning techniques, particular recursive neural networks, can also be effective [6], [7]. This work will focus on designing a BMI decoder using a spiking neural networks (SNN).

SNNs are a category of neural network that take trains of events, or spikes, as inputs, rather than real numbers. SNNs can be modeled as connections of units called neurons, which integrate incoming signals and emit outgoing spikes in ways that closely model biological neurons [9]. These networks can be implemented as electronic circuits on mixed-signal analog processors known as neuromorphic chips. Due to the sparse nature of their inputs, SNNs implemented on neuromorphic hardware have been shown to be extremely power efficient [10]. This property is highly relevant in the field of BMI development, since one of the major bottlenecks for intracortical BMI is power consumption. In order to avoid damaging neural tissue, BMI components must dissipate less than 40 mW/cm^2^ [11]. Unlike their counterparts, SNNs are highly power efficient, consuming as little as 4 *μ*W per neuron with an average firing rate of 40 Hz [12].

Due to their high energy efficiency and close resemblance to biological systems, there have been attempts to use SNNs for BMI decoders. One study developed a SNN decoder that replicates the behavior of a Kalman filter [11], while another used Spike Time Dependent Plasticity, a popular unsupervised learning method, to train a SNN-based BMI [13]. While these methods remain popular, recent research has shown that the accuracy of SNNs can be far improved using error backpropagation [14]. One recent work used a SNN with backpropagation to perform neural decoding with comparable accuracy to a state-of-the-art neural network [15]. However, backpropagation methods are less biologically plausible than STDP, and require significant memory overhead. No work has been done yet to evaluate the performance of biologically plausible backpropagation-based SNNs for neural decoding. Additionally, research on the use of SNNs for decoding in multiple regions of the brain is extremely limited, with most studies focusing only on the premotor cortex.

In this work, we design an SNN for decoding neural data and train the network parameters using a biologically plausible local learning rule. This study uses the Leaky Integrate and Fire neuron model to imitate the behavior of a neuron. The model is trained and evaluated two data sets, containing neural data from the premotor cortex of macaque primates [16] and from the hippocampus of rats [17], respectively. The Pearson Correlation Coefficient (*ρ*) is used to evaluate the accuracy of the model. To benchmark the model’s performance, we compare it to the accuracy of a Kalman Filter and Long Short-Term Memory artificial neural network (LSTM) when trained on the same data sets.

## II. Data Collection and Neural Processing

This work evaluates the SNN on two different data sets, each containing neural and kinematic data. The first study includes extracellular neural recordings from macaque primates [16], [18]. The study recorded neural activity from the premotor cortex of the monkeys as they performed reaching tasks. For each reaching task, the monkey moved a cursor towards an indicated target on a 20 cm by 20 cm computer screen. A total of three recording sessions from one monkey are present, for a total of approximately 33 minutes of data. This data set also contained kinematic data from each of the reaching tasks. The velocity of the on-screen cursor was recorded at 1 kHz, then downsampled and placed in 10 ms bins.

The second study included neural recordings and kinematic data from male Long-Evans rats [17], [19]. The study recorded neural activity from the right dorsal hippocampus of the rats as they pursued drops of water or food on a 180 cm by 180 cm platform. This paper will use 3 recording sessions containing a total of 233 minutes of data. The data set also recorded the position of the rat, which was sampled at 39.4 Hz.

For both studies, spikes were extracted from raw neural data using threshold crossing, and then sorted into independent neural units. The spiking data was placed in bins to pass as input to the SNN, with 10 ms bins used for the premotor cortex data and 20 ms bins used for the hippocampus data. For the premotor cortex data, independent reaching tasks were passed to the SNN as separate training examples, while the hippocampus data was split into 10 second long segments to provide multiple training examples for the SNN.

## III. Spiking Neural Network Implementation

### A. Neuron and Synapses

Spiking neural networks are very intuitive to apply to the problem of neural processing because they seek to mimic the behavior of biological neurons. SNNs take as input a sequence of discrete events known as a spike train, defined formally as *S*(*t*) = ∑*_keC_δ*(*t* – *t*^(*k*)^), where C is the set of all events and t^(*k*)^ represents the timing of event k in C [20]. Neuron units in a given layer l in an SNN perform operations on weighted sums of all the input spike trains to layer l, then emit their own spike trains, which are passed as input to the layer l+1. As a result, all information is propagated in the form of spike trains. This very closely resembles biological neurons, which receive sequences of sudden, discrete spikes from other neurons via their dendrites, and fire spikes in voltage known as action potentials.

For this study, we use the leaky integrate-and-fire (LIF) model to control the behavior of neuron units [9]. The LIF neuron maintains two internal values: the synpatic current, I(t), and the membrane potential, U(t). The synaptic current and membrane potential of the i-th neuron in a layer L in the network are governed by following:

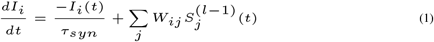

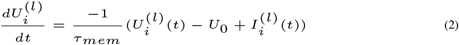

where W represents the matrix of weights applied to the input spikes and *τ_syn_* and *τ_mem_* are time constants [21]. To determine when the neuron emits a spike, the Heaviside step function is applied to the neuron’s membrane potential, with the spike train for the i-th neuron denoted by 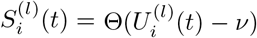. Here *ν* is a constant threshold value. After a neuron fires a spike, its membrane potential is reset to the resting potential, *U*_0_. This spike train is then passed as input to the next layer of neurons.

### B. SNN Architecture

The SNN used in this study consists of an input layer, with size corresponding to the number of neural input units from each recording session, two hidden layers of 65 and 40 neurons, respectively, and a readout layer. The sizes of the hidden layers were optimized using Bayesian Optimization [22]. The readout layer consists of 2 neurons that each output a continuous time signal, representing the predicted x and y components of the velocity of the primate’s hand.

### C. Regression Learning Methods

Spiking neural networks are typically applied for classification tasks, in which the readout layer consists of several spiking neurons and the output is determined by selecting the neuron with the highest spiking rate [14]. This method would not work for performing a regression task. Instead, the neural network must produce two functions in time, for the x and y components of the position of velocity data.

In order to best match the nature of the data from each data set, we developed two different methods for generating an output. The first method is to use two modified spiking neurons as the output layer of the function. These modified spiking neurons still maintain a synaptic current and membrane potential, but they never emit spikes, so the membrane potential, U(t), is never reset. The output of the model is simply the membrane potential. An advantage of this method is that the output is both continuous and time-differentiable, two properties which are shared by the actual kinematic data. Additionally, when given an input of duration T timebins, an output of equal duration is produced. However, a disadvantage of this method is that the membrane potential always starts at zero, which can lead to a poor fit when the kinematic data starts far from zero. A similar output encoding method was used in [15], but our method was developed independently prior to the release of that work.

The second output method uses a simple linear weighted sum of the outputs of the last hidden layer for multiple time steps. All hidden layers still consist of LIF spiking neurons, so the benefits of spiking neurons still apply. This output encoding method can start at any value, since the output does not depend on integrating any internal value, but it is also not continuous, and can be prone to noise.

We determined that the membrane potential encoding method was preferred for arm velocity decoding, while the rate encoding method was preferred for positional data. Since the velocity data was recorded from sequential reaching tasks, the initial value was typically relatively close to zero, so the error from the initialization of U(t) was minimized. Additionally, the neural data and kinematic data from that data set were recorded at the same frequency, so it was beneficial to use a method that produced an equal duration output. For the positional data, it was common for the rats to start far from the center of the platform, so it was necessary to use an output method that did not start at 0. Additionally, the kinematic data was recorded at a much lower frequency than the neural data, so it was beneficial to use a method where the duration of the output could be controlled.

### D. Loss Function and Learning Rules

To train the network, the loss function is defined as the sum of the mean squared errors of the actual and predicted x and y velocities. The SNN is trained using a local learning rule, in which a readout layer is appended to each hidden layer in the network, and the global loss is defined as the sum of the losses for each readout layer. Local learning requires significantly less memory overhead when training the network compared to a network that uses full error backpropagation. Additionally, it is considered more biologically plausible than standard backpropagation, since it does not require information about other neurons in order to update the weights for an individual neuron [23]. The synaptic weights were updated using surrogate gradient descent, a modified version of gradient descent in which the heaviside function, 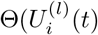, is replaced with a continuous, differentiable function 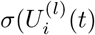 to make it possible to compute a gradient over a sequence of spikes [21].

## IV. Experiment and Results

The SNN described in the previous section was trained and tested on both data sets. Bayesian optimization was used to optimize various hyper-parameters, including hidden layer sizes, synapse and membrane time constants, learning rate, and dropout rate [22]. The SNN was trained for a total of 100 epochs, after which point it was clear that learning had stagnated. Each session was split into 3 subsets, with 60% of the reaches used for training, 20% for validation, and 20% for testing. The SNN achieved peak correlations of *ρ* = 0.77 for the premotor cortex data and *ρ* = 0.80 for the hippocampus data. The same data sets were also evaluated using a Kalman filter from the neural decoding tool set [24], as well as a backpropagation-based SNN and a LSTM network to benchmark the SNNs performance. As shown in Fig. 3 above, the SNN achieved a higher correlation than the Kalman filter and a lower correlation than the LSTM on the premotor cortex data, while all three methods performed equally on the hippocampus data. Across both data sets, there was no significant difference in accuracy between the local learning and backpropagation based SNNs. Additionally, a visual representation of the SNN performance can be seen in Fig. 4, which shows a samples of the predicted kinematic data alongside the measured kinematic values from both data sets.

**Fig. 1.**
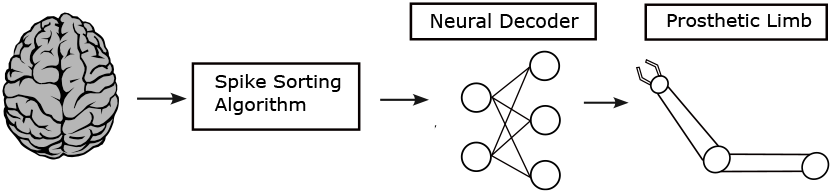
An illustration that gives a basic overview of the different components of a brain machine interface. In this process, raw neural data is taken from the brain, sorted into spikes, and then processed by a neural decoder to determine kinematic data. The kinematic data is then sent to a prosthetic. This work will focus on an implementation for the neural decoder as a spiking neural network. (Brain image taken from [8] under universal creative commons license.)

**Fig. 2.**
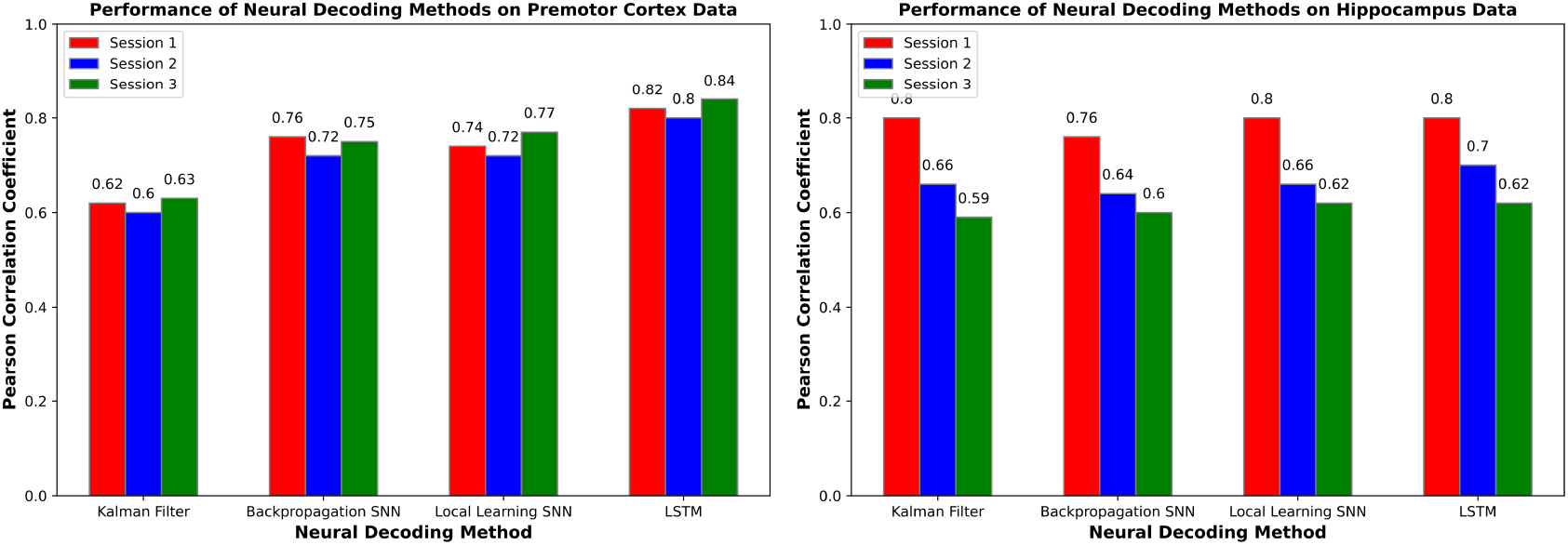
A graphical depiction of the spiking neural network architecture described below. The study used Bayesian optimization to obtain the optimal hyper-parameters. Specifically, the study optimizes the number of layers, number of neurons in a layers, drop-out rate, and regularization.

**Fig. 3.**
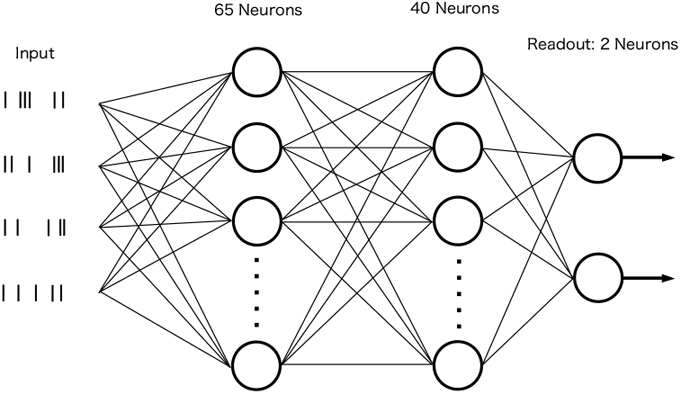
Bar plots showing the average performance of the Kalman Filter, SNN, and LSTM network on data from the three recording sessions. The correlations in the x and y coordinates were averaged. The SNN was evaluated using both backpropagation and local learning rules.

**Fig. 4.**
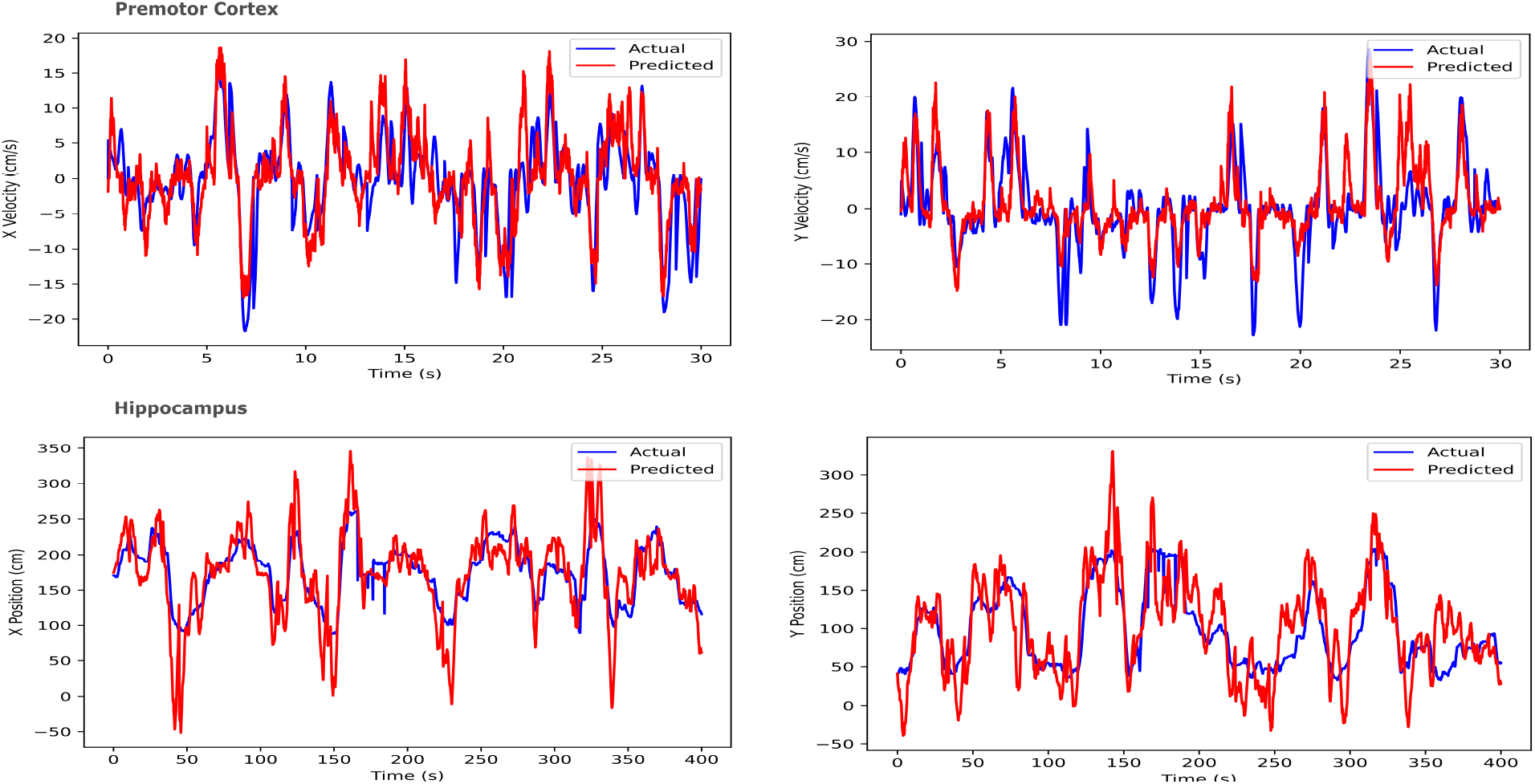
Two samples comparing the recorded kinematic data with the predicted kinematic data from the SNN output. The top graphs represent a 30 second sample taken from the premotor cortex data set, while the bottom ones represent a 400 second sample from the hippocampus data set. The x components of the kinematic data are on the left, while the y components are on the left.

## V. Discussion

This work demonstrates a biologically plausible spiking neural network for use in a brain machine interface system. Specifically, the SNN performs a neural decoding task on neural and kinematic data taken from monkeys and rats, determining the hand velocity of the monkeys and spacial position of the rats using data from the premotor cortex and hippocampus, respectively. The SNN uses the LIF model to control the behavior of its neurons, and trains using a local learning rule with surrogate gradient descent and is evaluated by finding the correlation between the output of the network and the true kinematic data. The results of this study show that the SNN achieves comparable performance or superior performance to a standard Kalman Filter neural decoder and a LSTM recurrent neural network, with the potential for significantly reduced power costs compared to these methods. Additionally, the study demonstrates that a SNN with a biologically plausible local learning rule achieves equal accuracy compared to a SNN using backpropagation. Future work will focus on designing and testing a hardware implementation of the SNN using analog circuitry. One of the main reasons for using a SNN in a brain machine interface is the comparatively low power needed to run a SNN.

